# Genome sequence of *Jaltomata* addresses rapid reproductive trait evolution and enhances comparative genomics in the hyper-diverse Solanaceae

**DOI:** 10.1101/335117

**Authors:** Meng Wu, Jamie L. Kostyun, Leonie C. Moyle

**Affiliations:** Department of Biology, Indiana University, Bloomington, Indiana, USA; Department of Plant Biology, University of Vermont, Burlington, Vermont, USA

**Keywords:** comparative genomics, de novo assembly, floral evolution, potato, tomato, transposable element

## Abstract

Within the economically important plant family Solanaceae, *Jaltomata* is a rapidly evolving genus that has extensive diversity in flower size and shape, as well as fruit and nectar color, among its ∼80 species. Here we report the whole-genome sequencing, assembly, and annotation, of one representative species (*Jaltomata sinuosa*) from this genus. Combining PacBio long-reads (25X) and Illumina short-reads (148X) achieved an assembly of approximately 1.45 Gb, spanning ∼96% of the estimated genome. 96% of curated single-copy orthologs in plants were detected in the assembly, supporting a high level of completeness of the genome. Similar to other Solanaceous species, repetitive elements made up a large fraction (∼80%) of the genome, with the most recently active element, *Gypsy*, expanding across the genome in the last 1-2 million years.

Computational gene prediction, in conjunction with a merged transcriptome dataset from 11 tissues, identified 34725 protein-coding genes. Comparative phylogenetic analyses with six other sequenced Solanaceae species determined that *Jaltomata* is most likely sister to *Solanum*, although a large fraction of gene trees supported a conflicting bipartition consistent with substantial introgression between *Jaltomata* and *Capsicum* after these species split. We also identified gene family dynamics specific to *Jaltomata*, including expansion of gene families potentially involved in novel reproductive trait development, and loss of gene families that accompanied the loss of self-incompatibility. This high-quality genome will facilitate studies of phenotypic diversification in this rapidly radiating group, and provide a new point of comparison for broader analyses of genomic evolution across the Solanaceae.

## INTRODUCTION

Understanding the genetic substrate of trait diversification is a longstanding goal in evolutionary biology. Diversification can involve a range of genetic changes, including point mutations in coding or regulatory regions, or structural variation such as chromosomal inversions or gene duplications (Stapley et al. 2010; Berner and Salzburger 2015). Until recently these data have been challenging to generate for all but the best-developed model species. However, the emergence of next-generation and single molecular sequencing technologies now allows the rapid generation of diverse genomic resources, including whole-genome sequences, transcriptome sequences and genome-wide marker panels for a much broader range of taxa (Stapley et al. 2010; Ellegren 2014; Bleidorn 2016). Comparative genomic analyses of related organisms provide opportunities to quantify species differences in genome size, complexity, noncoding features, and molecular evolution within genic regions, as well as structural differences in the number and identity of members of specific gene families, or in classes of transposable elements (TEs). In combination with data on specific phenotypic and functional trait variation, comparative genomic analyses can also evaluate the role of different genomic changes in both general patterns of lineage diversification and lineage-specific adaptive evolution. In addition to deciphering mechanisms of genome evolution, these data can be used to address the genetics of phenotypic diversity associated with adaptation and speciation, across groups of closely related, ecological diverse, species (Brawand et al. 2014; Zhang et al. 2014; Novikova et al. 2016; Yin et al. 2018).

The plant genus *Jaltomata* is one such rapidly radiating clade (Mione 1992; Miller et al. 2011). Consisting of 60-80 species, *Jaltomata* is closely related to both *Solanum* (which includes tomato, potato, and eggplant) and *Capsicum* (peppers), and together these three form a clade that is sister to the rest of the Solanaceae, a highly diverse plant family that also contains other economically important genera such as *Nicotiana* (tobacco) and *Petunia* (Bohs and Olmstead 1997; Olmstead et al. 1999; Walsh and Hoot 2001; Olmstead et al. 2008; Särkinen et al. 2013). Because multiple Solanaceous species already have whole-genome sequences with gene annotations, an additional high quality genome in the key phylogenetic position held by *Jaltomata* provides a valuable resource for clarifying the evolutionary relationships among these important clades, and for comparative analyses of genomic, genetic, and phenotypic evolution across the family. For example, although most Solanaceae species have the same base number of chromosomes (N=12; *Petunia* is the exception, with N=7), estimated genome sizes vary drastically (e.g. fourfold genome size difference between tomato and hot pepper) (Kim et al. 2014; Xu et al. 2017) suggesting that repetitive element divergence might be a key contributor to genome size variation. Further, comparative analyses of gene family evolution and sequence divergence could identify changes associated with important ecological trait variation (e.g. identification of tandemly duplicated genes involved in the capsaicin biosynthesis pathway specifically in hot pepper, (Kim et al. 2014; Qin et al. 2014). Comparative genomic analysis including *Jaltomata* therefore could reveal both large and small-scale genetic changes responsible for genomic, phenotypic, and functional differentiation among these important clades.

Apart from its key position within the Solanaceae, the genus *Jaltomata* itself varies widely in ecological range, vegetative characters, physiology, and reproductive form and function (Haak et al. 2014), making it a valuable emerging model for studies of adaptive diversification and evolution of novel traits. Most strikingly among its close relatives, *Jaltomata* lineages exhibit a unique diversity of derived floral traits, including in corolla (petal) shapes—that include rotate, campanulate, and tubular forms (Miller et al. 2011; Kostyun and Moyle 2017)—and in the amount and color of nectar produced, which ranges from essentially colorless to deep red (Hansen et al. 2007). In comparison, close relatives *Solanum* and *Capsicum* predominantly have flatter rotate corollas, and either colorless to pale yellow nectar (*Capsicum*) or no floral nectar at all (*Solanum*) (Knapp et al. 2004). *Jaltomata* species also vary in mature fruit color, including species that have either purple, red, orange, or green fruit at maturity; this fruit color variation appears to characterize three major subclades within the genus as separate dark purple, red, and orange-fruited clades (Miller et al. 2011; Särkinen et al. 2013; Wu et al. 2018). The orange-fruited clade is the subgroup containing most novel derived floral trait variation, and is estimated to have diverged within the last 1.5 million years, consistent with a very rapid recent radiation of floral and reproductive diversity that likely drew upon multiple sources of genetic variation (Wu et al. 2018). *Jaltomata* is also distinctive among Solanaceae genera in that all examined species are self-compatible (Mione 1992) (J. L. Kostyun and T. Mione, unpubl. data), whereas most other genera exhibit genetically determined self-incompatibility in some or all species (Goldberg et al. 2010). The availability of a high-quality genome in this genus could thus help to assess genome features specific to *Jaltomata* to identify genetic changes that might accompany or drive its unique and rapid trait evolution.

In this study, we generated a high-coverage and almost complete genome of one *Jaltomata* species, *J. sinuosa*, by adopting a hybrid assembly strategy using PacBio long reads and Illumina short reads. Using this newly assembly genome, we performed comparative genomic analyses with six additional high-quality genomes in the Solanaceae. We found that different topologies of *Jaltomata, Solanum*, and *Capsicum* were supported by a large number of individual gene trees, suggesting a complex history of rapid divergence and hybridization in the common ancestors of these three genera. Within *Jaltomata*, we identified a recent expansion of *Gypsy*-like elements around 1-2 million years ago (MYA) that likely contributed to the genome size expansion of *Jaltomata*. In addition, assessing genome features specific to *Jaltomata* identified genetic changes that could have contributed to rapid trait evolution in the genus, including loci with lineage-specific patterns of adaptive evolution, the loss of gene families that accompanied the loss of self-incompatibility, and the expansion of gene families potentially involved in novel trait development in *Jaltomata*. We discuss the significance of this genome assembly in the light of phenotypic diversity in this rapidly radiating group, and genomic evolution across the economically important plant family Solanaceae.

## MATERIALS AND METHODS

### Species selection and tissue sampling

Like its close relatives in *Solanum* and *Capsicum, Jaltomata* has its highest diversity in the Andean region of South America, although some species’ ranges extend into Central America and the southwestern United States. The 60-80 species within the genus are also distributed over a wide range of habitats —including tropical forests, coastal lowlands, and lomas (discrete fog-misted communities surrounded by arid desert)(Mione 1992; Mione and Yacher 2005). Of these, *Jaltomata sinuosa* was chosen for whole-genome sequencing because estimated heterozygosity within this species is the lowest of all evaluated species (Wu et al. 2018, and this paper), potentially facilitating genome assembly. Further, although *J. sinuosa* itself has rotate corollas with light-yellow nectar (i.e. similar to ancestral trait states in the genus), it belongs within the recently diverged and highly diverse orange-fruited clade, which incorporates the majority of novel floral diversity within the genus. To obtain genomic DNA, young leaf tissue was collected from a single individual of *J. sinuosa* grown in the Indiana University greenhouse (voucher available at IND herbarium), and flash frozen with liquid nitrogen. DNA was extracted using Qiagen DNeasy Plant Kits, purified with ethanol precipitation, and quality checked using Nanodrop and gel electrophoresis. Approximately 60µg genomic DNA was provided to the Duke University Sequencing and Genome Technologies facility for library preparation and sequencing: 4 large-insert (15-20kb) libraries were constructed and sequenced in 67 SMRT cells on the Pacific Biosciences (PacBio) platform.

### Genome assembly

We adopted a hybrid assembly approach (Koren et al. 2012) in which relatively high-accuracy Illumina paired-end reads (148X) were used to trim and correct low base-call-accuracy PacBio long-reads (25X). Initially, we used two different genome assembly strategies DBG2OLC (Ye et al. 2016) and MaSuRCA v3.2.2 (Zimin et al. 2017), and then evaluated the completeness of genome assembly for each using 1,515 plant near-universal single-copy ortholog within BUSCO v3 (Simão et al. 2015). We found the assembly output from MaSuRCA was much better than that from DBG2OLC, in terms of assembly coverage, contig N50, and genome completeness (Table S1). Thus, the initial assembly from MaSuRCA was used for all further analyses. Genome size was also estimated within the MaSuRCA pipeline based on the *k*-mer abundance distribution.

To remove potential contaminants in our assembly, all scaffolds were aligned against the NCBI non-redundant nucleotide sequence database using BLASTN with a cutoff of E-5. Scaffolds were assigned to the closest reference based on the best-combined hit score, and any scaffolds assigned to a non-plant species were removed (Bolger et al. 2014). In total, 22 small scaffolds were filtered out, consisting of 240,347 bases, leaving 7667 scaffolds in the genome assembly. To estimate assembly accuracy at the nucleotide level, we estimated the discrepancy between the sequences from genome assembly to the high-quality Illumina reads (see Supplementary text).

### Repeat annotation

We followed the “Repeat Library Construction-Advanced” steps from the MAKER-P pipeline (Campbell et al. 2014) to generate a *Jaltomata*-specific repeat library. Briefly, MITEs (miniature inverted transposable elements) were detected by MITE-Hunter (Han and Wessler 2010); LTRs (Long terminal repeats) were constructed using LTRharvest (Ellinghaus et al. 2008) followed by LTR_retriever (Ou and Jiang 2017); and other repetitive sequences were identified using RepeatModeler (http://www.repeatmasker.org/RepeatModeler.html). RepeatMasker (Smit et al. 2015) was then used to mask repeat elements in the assembled genome by searching for all homologous repeats in the species-specific library. To compare the recent activity of LTR elements, we also applied the same approach (i.e. LTRharvest followed by LTR_retriever) to annotate the full-length LTR elements in four other genomes from *Solanum* and *Capsicum*, including *S. lycopersicum, S. tuberosum, S. pennellii*, and *C. annuum*.

### Gene structure annotation

We followed the MAKER-P pipeline (Campbell et al. 2014) to annotate gene models in the assembled genome using three classes of evidence: RNA-seq data, protein homology, and *ab initio* gene prediction (see Supplementary text). MAKER-P synthesized all information from these three different classes and produced final annotations with high evidence-based quality, requiring annotation evidence distance (AED, which measures the goodness of fit of an annotation to the RNA/protein-alignment evidence supporting it) score <0.6. To assign gene functions, we followed the pipeline AHRD (https://github.com/groupschoof/AHRD) to automatically select the most concise, informative, and precise function annotation (see Supplementary text). For each gene, the associated gene ontology (GO) annotation(s) were assigned according to the predicted protein domain(s) (http://www.geneontology.org/external2go/interpro2go). To investigate the putative chromosomal locations of predicted genes in the genome, we performed whole-genome synteny alignment against the tomato genome using Satsuma v3.1.0 (Grabherr et al. 2010), and recorded the genes unambiguously associated with one identified syntenic region in tomato.

### Inference of homologous and orthologous gene clusters

We downloaded the annotated gene/protein-coding sequences of six other diploid Solanaceae species (including *S. lycopersicum, S. tuberosum, Capsicum annuum, Nicotiana attenuata, Petunia axillaris*, and *P. inflata*) from the SolGenomics database (http://solgenomics.net) and used *Arabidopsis thaliana* as the outgroup species. The six Solanaceae species were chosen because they each have one well-assembled and annotated genome (i.e. contig/scaffold size > 10kb and gene completeness > 95%) with a comparable number of annotated genes (∼35,000 genes). Identification of homologous gene clusters was performed using orthoMCL v2.0.9 (Li et al. 2003) with the default options. We obtained 3,103 single-copy 1-to-1 orthologous clusters that each contained one sequence from all seven species for downstream phylogenetic analyses. The coding sequences (CDS) of those 1-to-1 orthologs were aligned using PRANK v.150803 (Löytynoja and Goldman 2005) with codons enforced. As a quality check on all multiple sequence alignments, we removed poorly aligned regions using a sliding window approach that masked any 15-bp window from alignment if it had more than six mismatches (not counting indels/gaps, which were masked to N) among all the investigated species. After this process, any alignment with more than 20% of its sequence masked was removed from the analysis.

### Phylogenetic analyses

We reconstructed evolutionary relationships between *Jaltomata* and the six other Solanaceous species for which we had whole-genome data. We used four different, but complementary, inference approaches to perform phylogenetic reconstruction: 1) maximum-likelihood (ML) applied to concatenated alignments (in RAxML v8.23; Stamatakis 2006); 2) consensus of gene trees (RAxML V8.23, with Majority Rule Extended (Salichos and Rokas 2013); 3) quartet-based gene tree reconciliation (using ASTRAL v4.10.9; Mirarab and Warnow 2015); and 4) Bayesian concordance of gene trees (using MrBayes v3.2 (Huelsenbeck and Ronquist 2011) followed by BUCKy v1.4.4 (Larget et al., 2010)) (see Supplementary text). Using these different tree reconstruction approaches allowed us to evaluate the extent to which they generated phylogenies that disagreed, as well as to identify the specific nodes and branches that were robust to all methods of phylogenetic reconstruction. In addition, we also reconstructed and annotated the chloroplast and mitochondrial genomes in *Jaltomata* and performed phylogenetic analyses of *J. sinuosa* with *S. lycopersicum, C. annuum*, and *N. attenuata* (the species for which cpDNA and mtDNA sequences were readily available) using the concatenated sequences of mitochondrial or chloroplast genes in RAxML v8.23 (see Supplementary text).

Our phylogenetic analyses indicated that gene trees supported two conflicting bipartitions among *Jaltomata, Solanum*, and *Capsicum* (i.e. “(*Jaltomata, Solanum*), *Capsicum*” or “(*Jaltomata, Capsicum*), *Solanum*”) at roughly equal frequencies (see Results). To differentiate which of these topologies most likely represented the initial pattern of lineage splitting (i.e. the ‘true’ species tree) rather than relatedness due to subsequent introgression among species, we compared the relative divergence times (node depths) among species, using sets of gene trees that each supported one of the two most conflicting bipartitions, with *Nicotiana attenuata* as the outgroup. Gene trees constructed from non-introgressed (initial branching order) sequences are expected to have deeper mean divergence times at the two internal nodes (T_1_ and T_2_; Figure 3) than those constructed from introgressed sequences, because introgression will reduce sequence divergence between the two lineages that have exchanged genes (Fontaine et al. 2015). Internal node depths (divergence times) for each gene tree that supported one of these two alternative bipartitions were calculated from biallelic informative sites; per gene estimates were used to calculate the genome-wide means of divergence times T_1_ and T_2_, to determine which of the two topologies had higher average divergence times (see further, Supplementary text).

### Gene family analyses

To investigate changes in gene family sizes, we investigated the families that are significantly rapidly evolving among the six Solanaceae species including *Jaltomata*. (In this analysis, *C. annuum* was excluded because of the complex relationships among *Jaltomata, Capsicum*, and *Solanum*; see Results.). We determined the significantly expanded or contracted gene families along each branch of phylogeny using the program CAFE v3.0 (Han et al. 2013) with *P*-value cutoff of 0.01. The input gene families were generated from the OrthoMCL program (Li et al. 2003). Phylogenetic relationships were based on the output from the RAxML analysis of the concatenated dataset of 3,103 coding-sequence alignments. Divergence times among different species were directly retrieved from previous estimates among Solanaceae species (Särkinen et al. 2013). In addition, we further investigated the expansion of a specific candidate gene within *Jaltomata, SEUSS*, by examining gene trees, primary mapped Illumina reads (i.e. using only the best alignment of multi-mapped reads), PacBio reads spanning more than one gene copy, and expression patterns supported by RNA-seq from 14 *Jaltomata* species examined in our previous study (Wu et al. 2018) (see Supplementary text).

### Positive selection analyses

To infer positively selected genes in the *Jaltomata* genome, we tested 3,103 single-copy 1-to-1 orthologous genes (excluding *C. annuum*) whose gene tree topologies were the same as the inferred species tree. For each investigated gene, we inferred putative adaptive evolution (i.e. *d*_*N*_*/d*_*S*_ > 1) using the branch-site (BS) model (model = 2 and NS sites = 2) in PAML v4.4 (Yang 2007) on the terminal branch leading to *J. sinuosa.* This inference uses a likelihood ratio test (LRT) to determine whether the alternative test model (fixed_omega = 0) is significantly better than the null model (fixed_omega = 1), and identifies putative positively selected genes as those with a LRT *P*-value < 0.01 and a false discovery rate (FDR) less than 0.2 (Benjamini and Hochberg 1995). Since multi-nucleotide mutations (MNMs) can cause false inferences of positive selection in the PAML BS test (Venkat et al. 2017), for all significant genes we further applied a more conservative branch-site model (BS+MNM) (Venkat et al. 2017) in which another parameter *δ* is incorporated to represent the relative instantaneous rate of double mutations to that of single mutations (Venkat et al. 2017). We then assessed how many and which BS-significant genes remained significant (*P* <0.01) in the BS+MNM test. A GO enrichment analyses was performed on these remaining putative selected genes using ONTOLOGIZER v2.0 with the parent-child analysis and a cutoff *P*-value of 0.01 (Bauer et al. 2008).

## RESULTS

### Genome assembly and annotation

The genome of *J. sinuosa* is estimated to be ∼1,512 MB based on *k*-mer frequencies, which is consistent with the estimated size from flow cytometry (∼1,650 MB; D.C. Haak, unpublished data). Using the MaSuRCA genome-assembly pipeline, we generated an assembly of ∼1,456 MB, in which 96.6% of 1,515 BUSCO universal single-copy orthologous genes could be found (Table 1). The assembly comprises 7,667 scaffolds, with a contig and scaffold N50 of 364.9 KB and 397.6 KB, respectively (Table 1). In the ∼1,389 MB of genomic regions covered by at least five reads, we detected only 68,203 sites consistently supported by Illumina reads that were different from the assembled reference genome (including mismatches and indels), suggesting a lower bound of base calling error rate of ∼0.005%. The upper bound of error rates was estimated to be 0.28% by counting the total discrepancies between all the primarily aligned reads and the assembled reference, although the error rate might be higher in genomic regions that are not well covered by Illumina data (Schmidt et al. 2017). Overall, the high coverage of the plant conserved single-copy genes, low base-calling error rate, and the high mapping rate of RNA-seq reads (96.2%, see below), indicate a high quality assembly for the *J. sinuosa* genome. We also assembled the chloroplast genome into a single contig with a total length of 156 KB (GC content of 37.91%) with 82 annotated genes; the mitochondrial genome was assembled into a single contig with a total length of 317 KB (GC content of 42.05%) and 21 annotated genes.

**Table 1.**
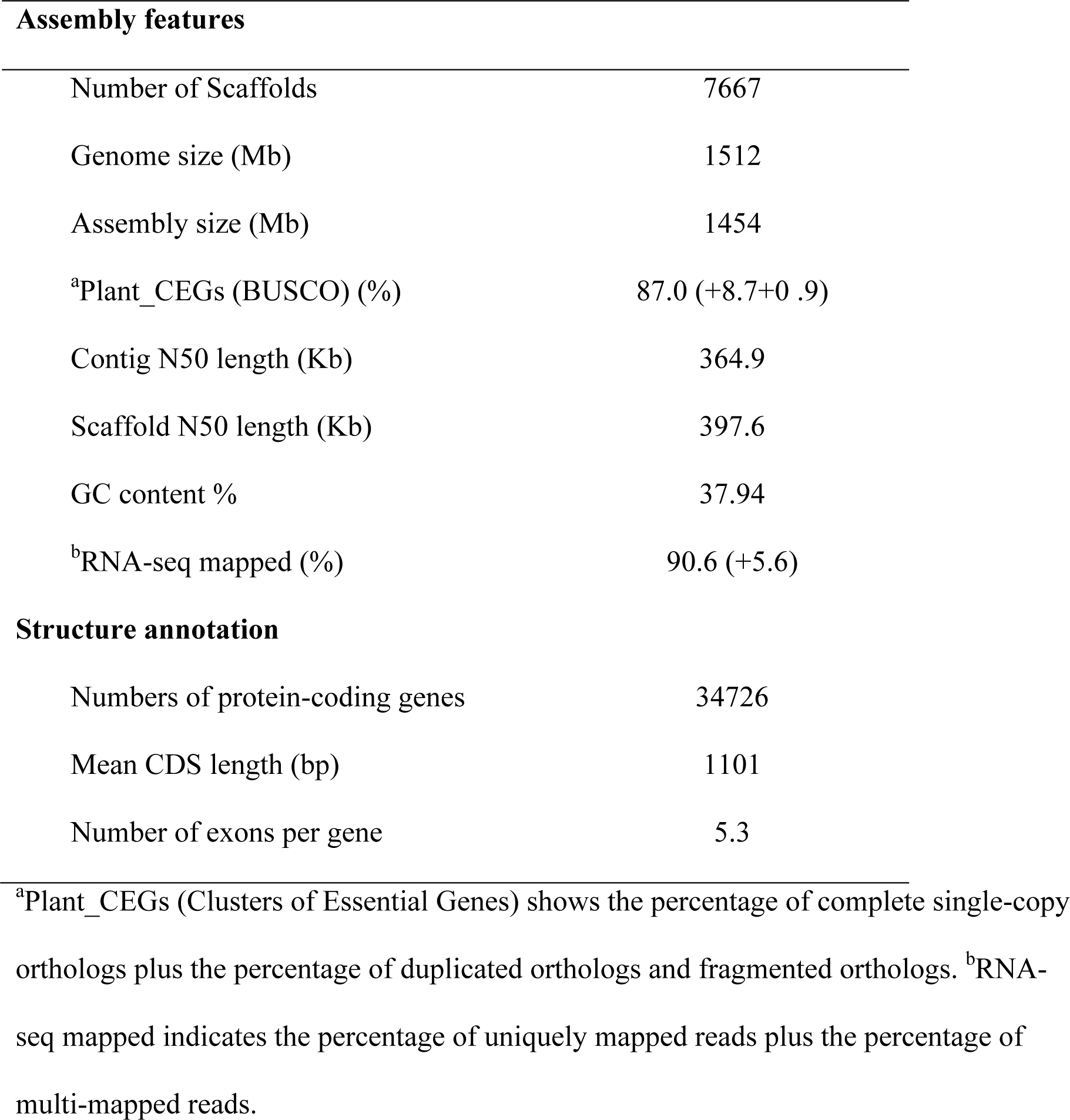
Summary of the *J. sinuosa* genome assembly.

| **Assembly features** |  |
| --- | --- |
| Number of Scaffolds | 7667 |
| Genome size (Mb) | 1512 |
| Assembly size (Mb) | 1454 |
| <sup>a</sup>Plant_CEGs (BUSCO) (%) | 87.0 (+8.7+0 .9) |
| Contig N50 length (Kb) | 364.9 |
| Scaffold N50 length (Kb) | 397.6 |
| GC content % | 37.94 |
| <sup>b</sup>RNA-seq mapped (%) | 90.6 (+5.6) |
| **Structure annotation** |  |
| Numbers of protein-coding genes | 34726 |
| Mean CDS length (bp) | 1101 |
| Number of exons per gene | 5.3 |

<sup>a</sup>Plant\_CEGs (Clusters of Essential Genes) shows the percentage of complete single-copy orthologs plus the percentage of duplicated orthologs and fragmented orthologs. <sup>b</sup>RNA-seq mapped indicates the percentage of uniquely mapped reads plus the percentage of multi-mapped reads.

### Repetitive element annotation and LTR insertion age distribution

The assembled *J. sinuosa* genome contains a total of ∼1,158 MB (80.29% of the assembly) of repetitive sequences (Table S2). LTR retrotransposons are the major source of repetitive sequences, accounting for 64.43% of the genome assembly (Figure 2A); of these, *Gypsy* elements are much more abundant (506.1 MB) than *Copia* elements (59.5 MB) (Figure 2A; Table S2). To evaluate the recent activity of LTR elements, we identified 1,682 full-length LTR retrotransposons (including 823 *Gypsy* and 151 *Copia*; Table S3) and, using the distribution of sequence divergence between the two terminal repeats within each LTR element, we infer a recent burst of *Gypsy* element activity around 1-2 MYA (Figure 2B). In comparison, the *C. annuum* and three *Solanum* genomes (*S. lycopersicum, S. tuberosum* and *S. pennellii*) show many fewer recent insertions of *Gypsy* elements (Figure S1B-D). Instead, *Gypsy* elements in *C. annuum* are inferred to have been most active around 3 MYA (Figure S1A), whereas a higher abundance of recently active *Copia* elements was detected in *S. pennellii* (Figure S1D), consistent with a previous study (Bolger et al. 2014).

**Figure 1.**
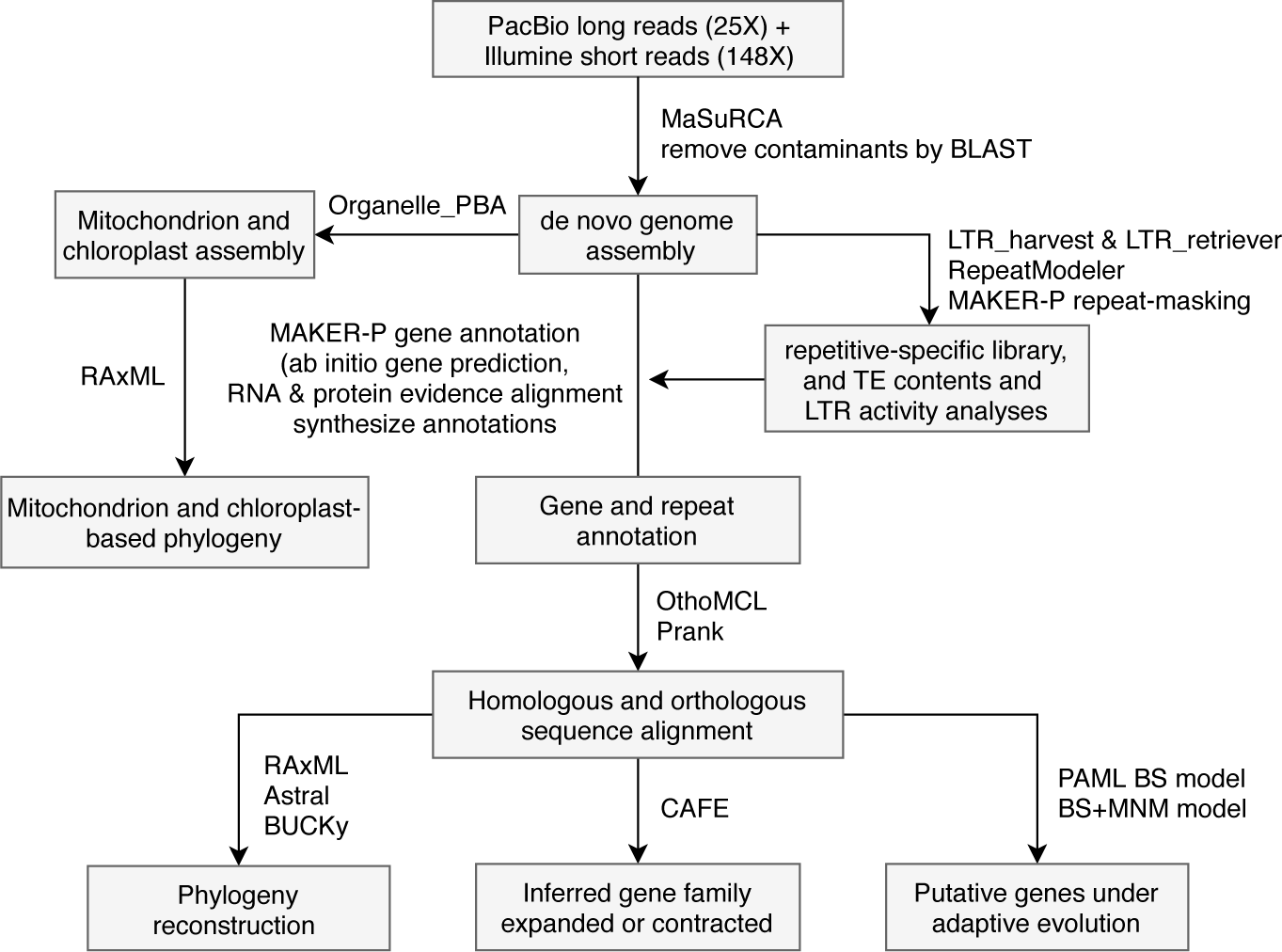
Workflow of assembly of *J. sinuosa* genome and downstream comparative genomic analyses

**Figure 2.**
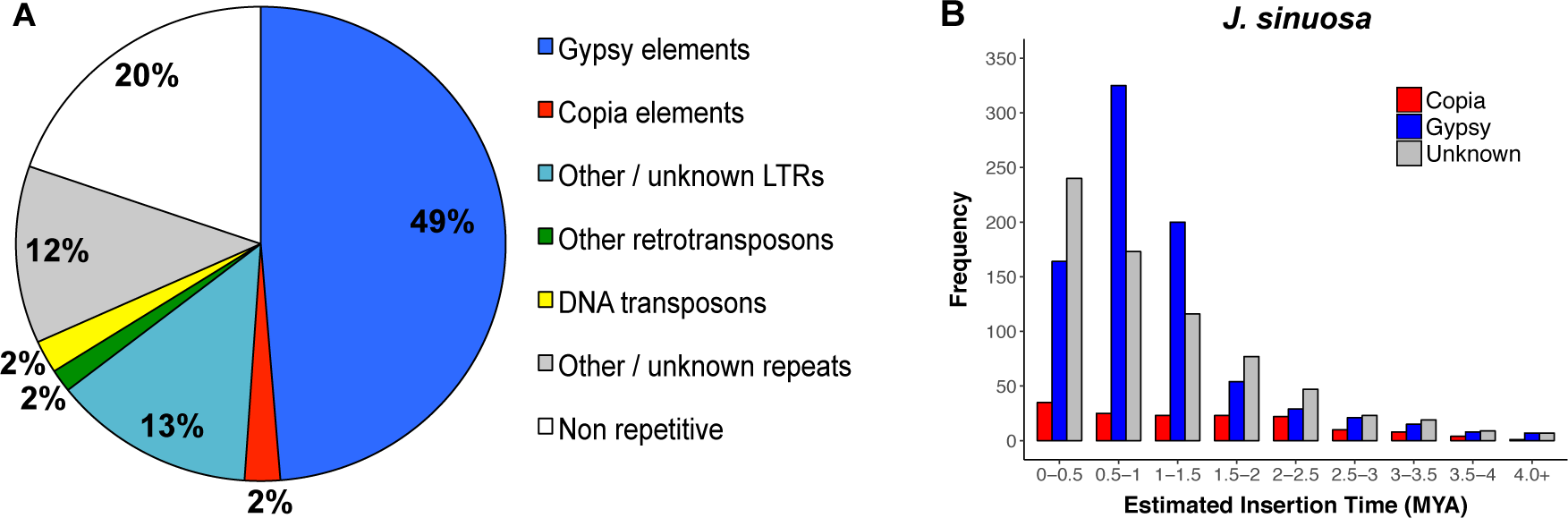
Landscape of repetitive sequences in the *J. sinuosa* genome. **(A)** The repeat contents in the assembly. **(B)** The estimated insertion age distribution of full-length LTR elements in the assembly.

### Gene annotation and transcription

A total of 34,726 high-confidence (AED <0.6) protein-coding genes with 37,106 transcripts were predicted (Table S4), which is similar to the 35,768 predicted genes in the domesticated tomato, *S. lycopersicum* (The Tomato Genome Consortium 2012). The protein coding sequences (CDS) in the annotated genes have an average length of 1,101 bp and the predicted genes have an average of 5.3 exons (Table S5); both are similar to the average CDS length (∼1,027 bps) and exon number (∼4.9) in the tomato annotation ITAG3.2 (The Tomato Genome Consortium 2012). Through whole-genome synteny alignment, 23,013 (66.3%) of these predicted *Jaltomata* genes were associated with unambiguous syntenic regions within the tomato genome (Table S6). Among all annotated genes, nearly all of them (99.90%) were functionally annotated through the AHRD pipeline (Table S7). The transcriptome-wide RNA-seq reads (Wu et al. 2018) from *J. sinuosa* were mapped against the assembly with a success rate of 96.2%, with 67.81% of annotated genes having more than 0.5 transcripts per million (TPM >0.5). Because RNA-seq data from one species might only sample a subset of expressed genes, we also mapped the RNA-seq reads from 13 other *Jaltomata* lineages (Wu et al. 2018) back to the assembly, and identified 82.83% of annotated genes that have TPM >0.5 in at least one species. We identified 11,563 gene families that were shared among *J. sinuosa, S. lycopersicum, C. annuum, N. attenuata* and *P. axillaris*, while a total of 953 gene families were specific to *J. sinuosa* (Figure 4A). GO term enrichment analysis indicated these *J. sinuosa*-specific genes are significantly over-represented in negative regulation of metabolic process or catalytic activity (Table S9).

### *Complex phylogenetic relationships between* Jaltomata, Solanum, *and* Capsicum

The four phylogeny-reconstruction methods all supported the same topology among the seven Solanaceae species, that included *J. sinuosa* as more closely related to *C. annuum* than to *S. lycopersicum* (Figure 3A; Figure S2). However, there was also substantial gene tree discordance observed at this node: only 42% of gene trees supported the branch that groups *J. sinuosa* together with *C. annuum*. An internode certainty (IC) value of zero on this branch indicates that the number of gene trees supporting this bipartition is almost equal to the number of gene trees supporting the most common conflicting bipartition (Figure 3A). A similar pattern was also observed at the internode that splits the two *Petunia* species from the other investigated Solanaceae species (Figure 3A).

**Figure 3.**
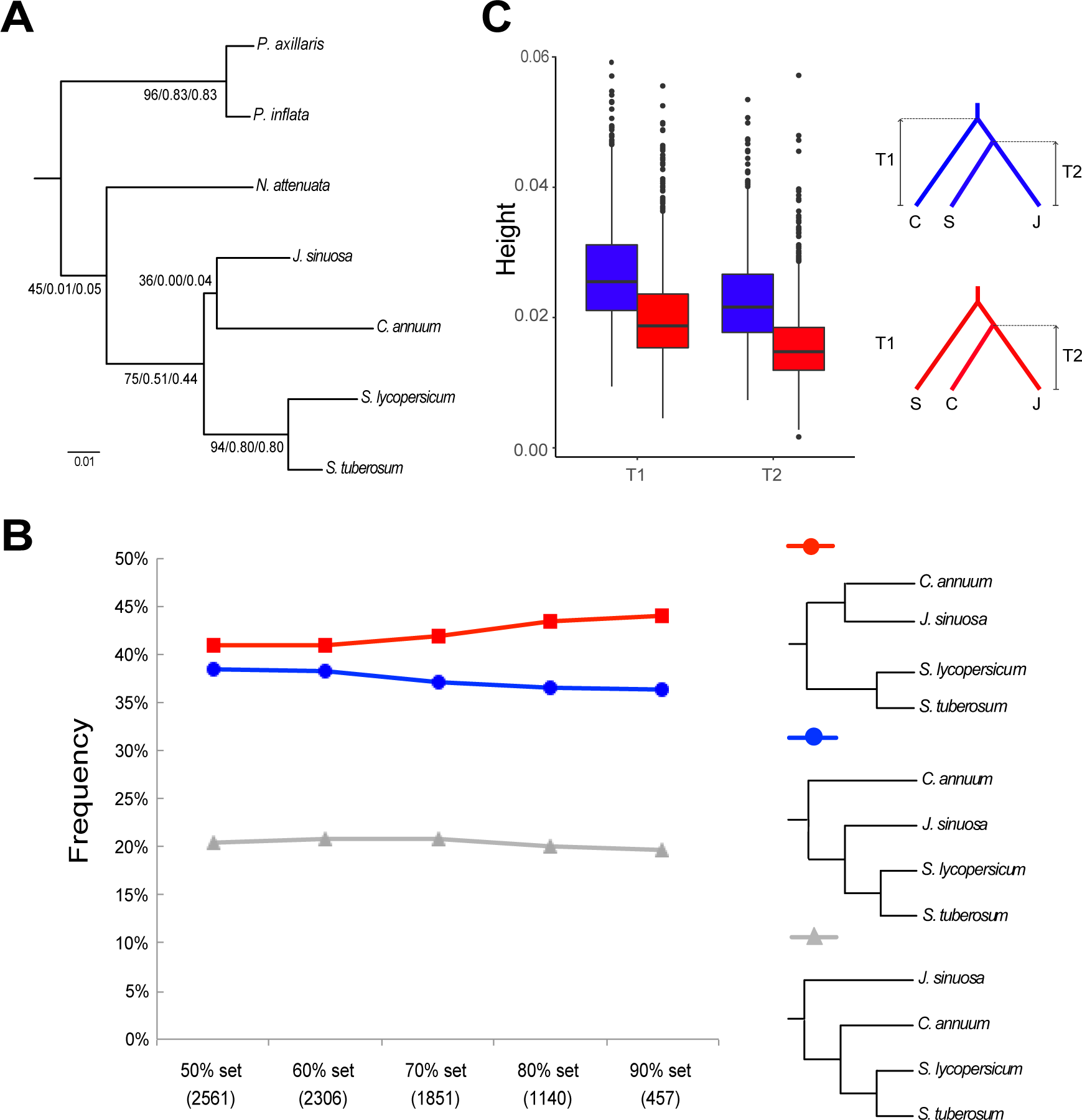
Phylogenetic relationships among the seven investigated Solanaceae species. **(A)** Concatenated RAxML tree rooted by *A. thaliana*. The values above each internal branch indicate the gene tree concordance factor (%), internode certainty (IC), and tree certainty all (TCA) supporting each node in the majority-rule consensus tree, from BUCKy (Larget et al. 2010). **(B)** The percentage of gene trees supporting the three alternative phylogenetic positions of *Jaltomata* relative to *Solanum lycopersicum, Solanum tuberosum* and *Capsicum annuum* (assuming *Solanum* is monophyletic). Five different sets of gene trees were used, which were selected based on the average bootstrap cutoff across the gene tree. (**C**) Evaluation of the relative node depths (ages) in the two alternative bipartitions among *Jaltomata* (J), *Solanum* (S) and *Capsicum* (C). The higher average height (depth) of T1 and T2 indicate that *Jaltomata* and *Solanum* are sister clades, whereas shallower gene trees supporting *Jaltomata* and *Capsicum* as sister are likely to be influenced by introgression.

In order to further investigate the phylogenetic placement of *Jaltomata* relative to *Solanum* and *Capsicum*, we examined the proportion of gene trees supporting the each of the three different possible topologies, using five groups of genes which had progressively higher bootstrap support (i.e., average % bootstrap support across the RAxML gene trees of greater than 50, 60, 70, 80, or 90%). For the largest (>50% bootstrap) group, we found that ∼42% of gene trees supported the most common bipartition (i.e. (*Jaltomata, Capsicum*), *Solanum*), but that ∼38% gene trees supported the conflicting topology that places *Jaltomata* as sister to *Solanum*; this general pattern is consistent across sets of genes with increasingly higher power (Figure 3B). Similarly, our phylogenies constructed from the concatenated mitochondrial and chloroplast gene sequences indicated that ten mitochondrial genes supported *Jaltomata* as the sister clade of *Capsicum*, while 72 concatenated chloroplast genes supported *Jaltomata* as closer to *Solanum* (Figure S3).

To differentiate which of the two majority conflicting topologies is due to the initial lineage splitting events versus subsequent introgression, we compared the estimated divergence times among the three species from those genes supporting either of the two conflicting topologies. The gene trees supporting “*Solanum*, (*Capsicum, Jaltomata*)” have lower mean divergence times T_1_ and T_2_ relative to those from the gene trees supporting “*Capsicum*, (*Jaltomata, Solanum*)” (Figure 3C). Because introgression reduces sequence divergence between species exchanging genes (see Methods), our data suggests that the initial species branching order was “*Capsicum*, (*Jaltomata, Solanum*)”, while the excess of gene trees supporting “*Solanum*, (*Capsicum, Jaltomata*)” is due to subsequent introgression between *Capsicum* and *Jaltomata*.

### *Dynamic evolution of gene families in* Jaltomata

We identified 129 rapidly evolving gene families that contracted specifically on the branch leading to *J. sinuosa* (Figure 4B). Interestingly, of these we found a gene family (cluster 7; Table S10) that is functionally involved in pollen-pistil interactions (GO:0048544) for which there are only 21 genes in the *J. sinuosa* genome, while the other five investigated Solanaceae genomes have 38-46 genes (Table S11). This contracted gene cluster included receptor-like kinase family proteins with an S-locus glycoprotein domain, which are involved in plant reproduction and signaling in pollen-pistil interactions (Table S10). Based on our analysis of syntenic blocks with the tomato genome, the genes in this cluster are distributed across multiple chromosomes in the genome (Table S10). The loss of multiple putative S-locus receptor kinase family proteins is consistent with the ancestral loss of self-incompatibility in all *Jaltomata* lineages (see Discussion). Compared to *Jaltomata*, contraction of different gene families was detected in the *S. lycopersicum* and *N. attenuata* genomes (Table S12).

**Figure 4.**
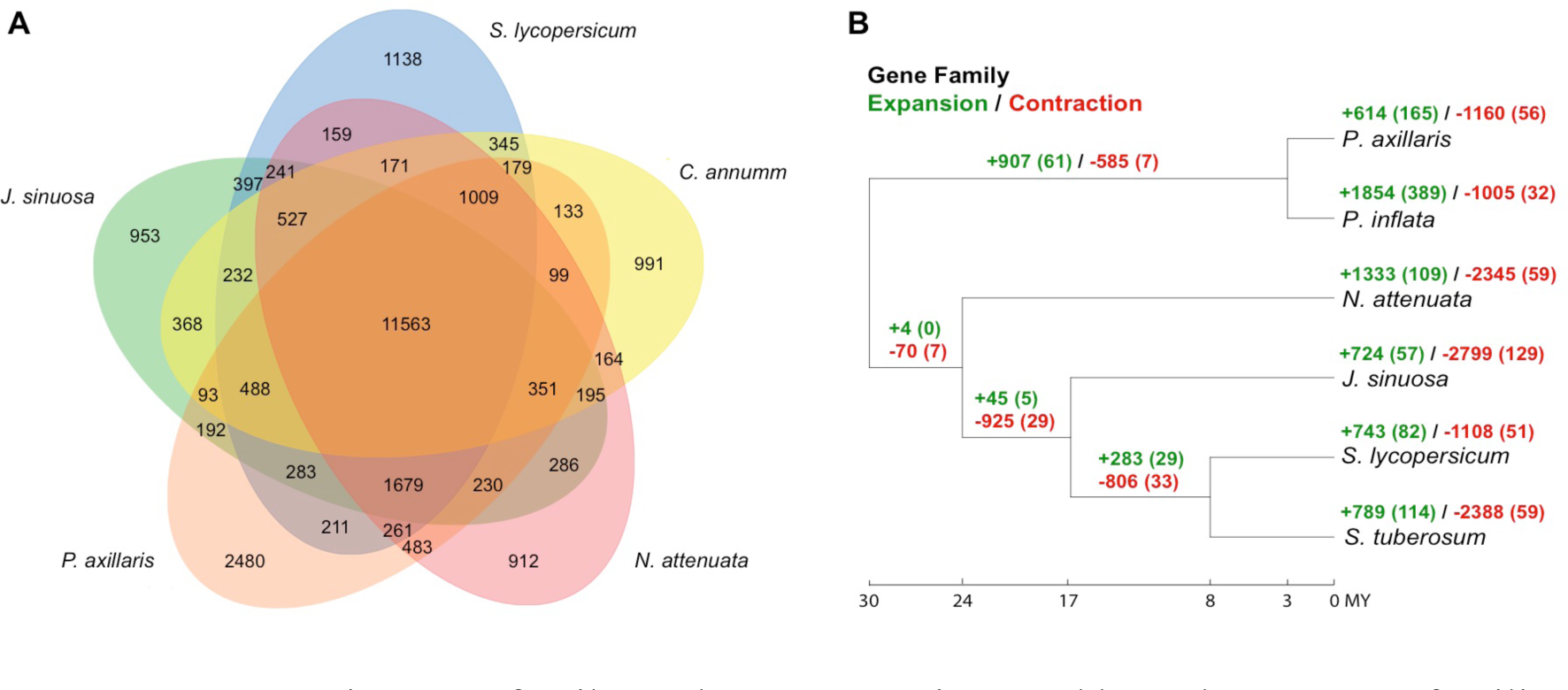
Comparative gene family analyses. **(A)** Unique and homologous gene families. The numbers of unique or shared gene families are shown in the corresponding diagram components. **(B)** Gene family expansion and contraction patterns across the Solanaceae. The numbers of expanded (green) and contracted (red) gene families are shown in each branch, and numbers in the brackets are the number of significantly expanded or contracted gene families (*P* <0.01).

We also identified 57 rapidly evolving gene families that expanded specifically on the branch leading to *J. sinuosa* (Figure 4B). These gene families were involved in a broad range of functions, including genes responsible for plant stress-related response, such as disease resistance, response to wounding, and regulation of nitrogen compound metabolic process (Table S13). Similar to *J. sinuosa*, significant lineage-specific expansion of various stress-related gene families was also found in other Solanaceae species, including genes involved in heat shock, oxidative stress and resistance to fungi in *S. lycopersicum* and *N. attenuata* genomes (Table S14). Of the expanded families specific to the *Jaltomata* genome, one particularly interesting example involved the transcription factor *SEUSS*, which is known to play an important role in floral organ development in model systems (see Discussion). While only one copy of *SEUSS* was identified in each other Solanaceae genome, an estimated ten copies (seven of them were annotated by the MAKER pipeline) were detected in the *J. sinuosa* genome (Figure 5A), and the gene tree of these copies indicates the expansion of *SEUSS* happened recently and specifically in the *Jaltomata* lineage (Figure 5A). The inferred ten copies of *Jaltomata SEUSS* gene were found on two scaffolds, suggesting the expansion of *SEUSS* gene copy number is due to recent tandem duplication (Figure 5B).

**Figure 5.**
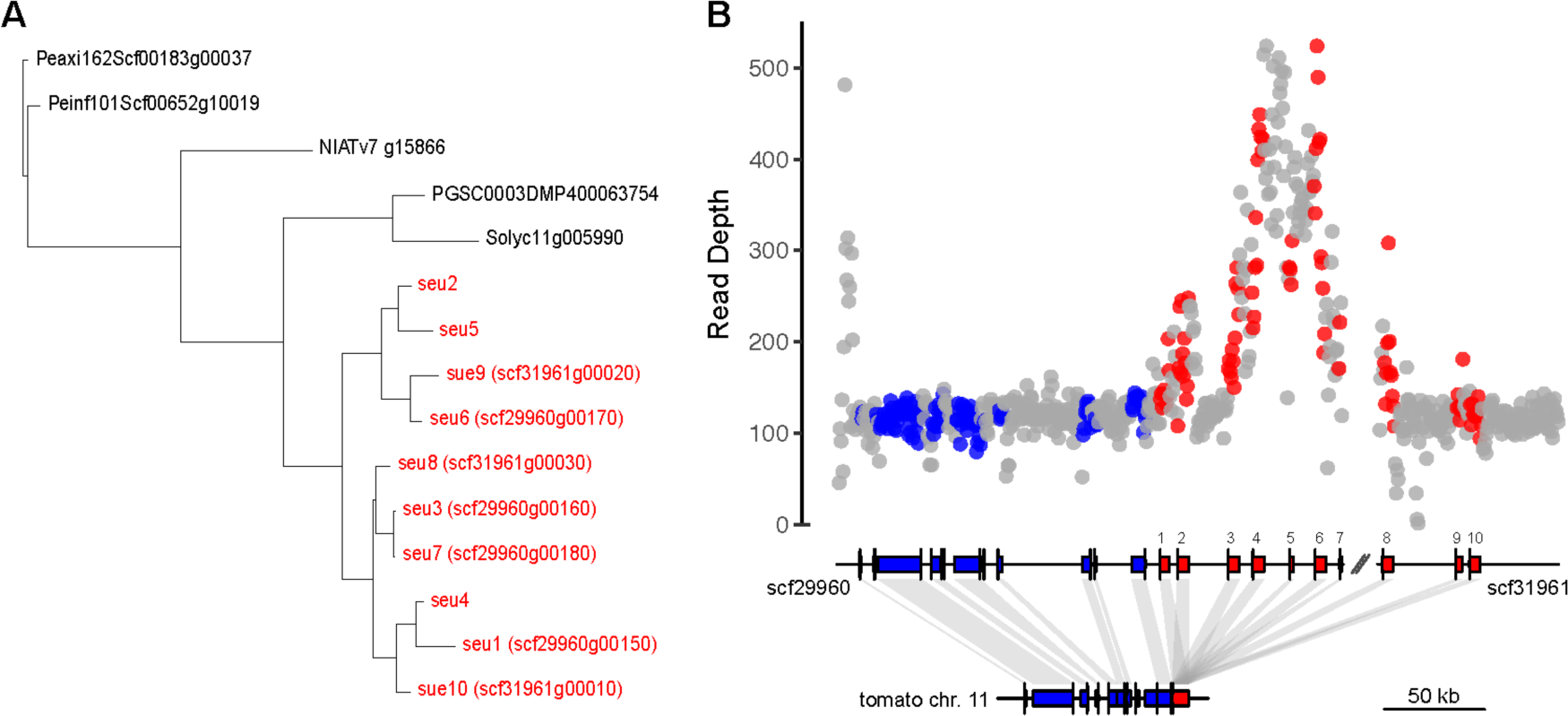
Evolutionary dynamics of copy number of the gene SEUSS in *J. sinuosa*. **(A)** Lineage-specific expansion of SEUSS in *J. sinuosa*. The 10 duplicated copies were labeled according to their relative position order along the located scaffolds. Among them, seven copies were annotated by the MAKER pipeline. **(B)** Validation that there are multiple copies of SEUSS (red) using read depth of Illumina DNA-seq reads which is at least equal or above the genomic background (grey) and adjacent single-copy genes (blue).

Several methods validated the presence of >1 *SEUSS* copies in *Jaltomata*. We identified five PacBio long reads spanning two adjacent *SEUSS* copies. We also found that read depth around each *SEUSS* copy was equal to or higher than read depth in the adjacent single-copy loci or background genomic regions (Figure 5B). The higher depth of primary mapped reads at some of these inferred *SEUSS* copies suggests that there might be additional paralogous copies of these loci that we were unable to differentiate because their sequences are too similar (i.e., due to very recent duplication events). Using RNA-seq data from (Wu et al. 2018), we found that four structurally intact copies (*SEUSS* 4, 6, 8, 10) have >25 reads that uniquely map to them in at least one of 13 *Jaltomata* species, suggesting there are at least four putatively functional copies in the genus (Table S15). Across all 13 species, only RNA-seq reads from the sampled reproductive, but not vegetative, tissues mapped to *SEUSS* loci (Table S15), and expression of *SUESS* copies varied among species: three copies (*SEUSS* 4, 6, 8) were expressed in all purple-fruited lineages (i.e. *J. repandidentata, J. procumbens, J*. *darcyana*) whereas almost all of the ten inferred *SEUSS* copies had no reads mapped (i.e. very low expression) in most of the orange/green-fruited lineages (including *J. sinuosa*) (Table S15). This difference in number of reads mapped in different *Jaltomata* lineages cannot be explained by sequence divergence, since the purple-fruited lineages are more distant from *J. sinuosa* relative to other investigated *Jaltomata* lineages.

### *Detection of genes potentially under positive selection in* Jaltomata

Using the branch-site model in PAML, we identified 89 genes out of 6,582 testable genes as putatively under positive selection (i.e. the lineage-specific selection model fit significantly better than the null model, *P*-value <0.01 and FDR <0.2; Table S16). After implementing the BS+MNM test (Venkat et al. 2017) on these 89 putative selected genes, 58 of them remained significant (*P*-value <0.01; Table S16). Some of these 58 loci are involved in stress responses, including resistance to osmotic and oxidative stress, response to heat shock, and heavy metal transportation (Table S16); however, these putatively selected genes were not enriched for any specific GO terms (Table S17). Nonetheless, among these loci are several candidates for elements of *Jaltomata*-specific trait evolution, including loss of self-incompatibility, growth during (floral) development, and regulation of pigment biosynthesis (see Discussion).

## DISCUSSION

Here we generated a high-quality genome sequence of a representative species (*J. sinuosa*) from within *Jaltomata*, a rapidly evolving, florally and reproductively diverse genus in the Solanaceae. We used these data to clarify a complex history of origin that *Jaltomata* shares with its two most closely related genera—*Solanum* and *Capsicum*—and to infer recent TE dynamics that might be responsible for genome-size evolution within the Solanaceae. We also identified overall patterns of significant gene family gain and loss, as well as adaptive molecular evolution, which could be implicated in the rapid reproductive trait evolution that is distinctive to *Jaltomata* among its Solanaceous relatives.

### *Comparative phylogenomic analysis reveals complex history of divergence among* Jaltomata *and its closest relatives*

The Solanaceae is a highly speciose plant family, with an estimated 100 genera and 2500 species that have all evolved within the last ∼30 MY. Previous phylogenetic studies have confirmed that many lineages within the Solanaceae arose within a highly compressed time frame (Olmstead et al. 1999; Särkinen et al. 2013), making resolution of some evolutionary relationships challenging. In particular, previous molecular phylogenetic studies using chloroplast and nuclear loci indicated that *Jaltomata* is close to both *Solanum* and *Capsicum*, however the relationship among these three genera has varied depending upon the specific loci used in phylogenetic reconstruction (Bohs and Olmstead 1997; Olmstead et al. 1999; Walsh and Hoot 2001; Olmstead et al. 2008; Särkinen et al. 2013). The *Jaltomata sinuosa* genome therefore provided an opportunity to evaluate and clarify the historical evolutionary relationships among key genera within Solanaceae.

Just as with other recent phylogenomic studies of contemporary (Brawand et al. 2014; Lamichhaney et al. 2015; Novikova et al. 2016; Pease et al. 2016) or more ancient rapid radiations (Jarvis et al. 2014; Wickett et al. 2014; Suh et al. 2015; Yang et al. 2015), we detected evidence for substantial gene tree discordance in relationships among the seven Solanaceous species analyzed here. This included high discordance (internode uncertainty) at the internode that split the *Petunia* lineages from the remaining species (Figure 3), as well as the internodes separating *Jaltomata, Solanum*, and *Capsicum*. In particular, our concatenation-ML phylogeny supported a closer relationship between *Jaltomata* and *Capsicum*, but we also detected a similar number of individual gene trees supporting the alternative topology of *Jaltomata* and *Solanum* as more closely related (Figure 3). The mitochondrial and chloroplast phylogenetic trees were also discordant with each other (Figure S3), with cpDNA supporting *Jaltomata* and *Solanum* as sister, whereas mitochondrial loci indicated that *Jaltomata* is sister to *Capsicum*. Genome-wide, the observed pattern of minority gene tree discordance was not consistent with the action of ILS alone (under ILS the two alternative minority trees are expected to be approximately equally represented; Degnan and Rosenberg 2009), indicating that a substantial component of discordance was likely also due to introgression between lineages (Huson et al. 2005).

Following the logical framework in Fontaine et al. (2016), in which the younger (shallower) tree topology is inferred to be due to introgression, we used the relative depth (age) of the two alternative tree topologies to infer that *Jaltomata* is likely sister to *Solanum* (a relationship supported by the tree with the older/deeper mean node depths). In contrast, an excess of gene trees supporting a sister relationship between *Jaltomata* and *Capsicum* (the tree with on average shorter/younger node depths) is likely to be due to introgression since the split of the three species; this is despite the observation that the latter tree is marginally more frequent across all gene trees compared to the next most common bipartition (Figure 3). Our use of whole-genome sequence data from *Jaltomata* therefore enabled us to disentangle the likely complex history of origin at the base of the clade that unites these three lineages.

Our analyses also highlighted the extensive phylogenetic incongruence among these and other Solanaceae genera more generally, a history that should be accounted for in comparative studies. In particular, assessing the level and distribution of incongruence is critical when making inferences about trait evolution in radiating lineages, as substantial gene tree discordance can contribute to incorrect inferences of convergence (‘hemiplasy’) at both the phenotypic and molecular level (Hahn and Nakhleh 2016; Wu et al. 2018).

### Transposable elements contribute to genome size evolution across the Solanaceae

We also used the *J. sinuosa* genome assembly to examine possible causes of genome size evolution, specifically variation in transposable element history. The estimated genome size of *J. sinuosa* (∼1.5 GB) is >50% larger than the tomato genome (∼0.9 GB), but less than half of the hot pepper genome (∼3.5 GB), despite no accompanying change in ploidy level, suggesting that transposable elements might be an important contributor to genome-size variation in the Solanaceae. Indeed, we determined that nearly 80% of the assembly consists of repetitive elements, with the vast majority belonging to the *Gypsy*-like family (Figure 2). We also inferred a recent proliferation of *Gypsy*-like elements that occurred around 1-2 MYA, which might have contributed to the larger genome size in *Jaltomata* lineages relative to *Solanum*. Previous studies have revealed a similar pattern of relatively recent *Gypsy* proliferation in other Solanaceae species, including a substantial excess of *Gypsy* elements (12 fold more frequent than *Copia* elements) within the hot pepper genome compared to domestic tomato (Kim et al. 2014). Similarly, genome size variation scales with variation in the number of *Gypsy* repeats among four *Nicotiana* species (Xu et al. 2017). In contrast, within *Solanum* the larger genome size of wild species *S. pennellii* compared to domesticated tomato is associated with a recent proliferation of *Copia*-like elements (Bolger et al. 2014). Overall, this and other recent analyses indicate that differential activity of LTR elements contributed substantially to differences in genome size across the Solanaceae, and suggest that future studies examining these and broader transposable element activity and DNA loss (Kapusta et al. 2017) could reveal important dynamics of genome size and content variation in this group.

### *Both gene family evolution and specific molecular changes contribute to unique reproductive trait variation within* Jaltomata

Unlike close relatives *Solanum* and *Capsicum*, species in *Jaltomata* exhibit extensive floral diversity in corolla shape and nectar volume and color (Miller et al. 2011) in addition to an apparently ancestral transition to self-compatibility (Mione 1992) (Kostyun and Mione, unpubl.). Another goal in developing a representative genome sequence for *Jaltomata* was therefore to identify genetic mechanisms that might have contributed to the evolutionary origin of these derived floral and reproductive states. In our previous phylogenomic study using transcriptome data, we investigated patterns of molecular evolution in several thousand loci across 14 species within *Jaltomata*, and found that relatively few adaptively evolving genes (i.e. with *d*_*N*_/*d*_*S*_ >1) were functionally associated with floral development. However, extensive gene tree discordance and very low sequence divergence (<1% among the species in the radiating group that displays the derived floral traits) only allowed a small subset of genes to be tested at some internal branches (Wu et al. 2018). Moreover this transcriptome-based study only focused on molecular variation in coding sequences, although regulatory changes and structural variation, such as gene duplications, are also known contributors to the evolution of novel traits (Hoekstra and Coyne 2007). The generation of a reference genome in this group therefore allowed us to address the possible role of additional genetic variation that could not be investigated previously.

### A. Contraction and expansion of specific gene families associated with reproductive trait evolution

In each of the seven Solanaceae lineages examined, we detected evidence for substantial expansion and contraction of gene families (Figure 4b), and several of these may have contributed to reproductive trait evolution specifically in *Jaltomata*. First, we detected significant contractions of gene families functionally associated with pollen-pistil interactions, including the putative S-locus receptor kinase family proteins, which could be a genomic signature of the ancestral transition to self-compatibility that appears to characterize this genus. Within the Solanaceae, self-incompatibility is mediated by the “S-locus” which encodes a single female/stylar S-determinant (i.e. S-RNase) and one or more pollen-expressed F-box proteins (Iwano and Takayama 2012) that interact to cause pollen rejection when the male and female parent share the same allele at the S-locus (McClure et al. 1989). Multiple studies have shown that loss of the pistil-side S-RNase mediates the transition from SI to SC in Solanaceous species, although the loss of one or more pollen-side F-box proteins and other components of the SI-machinery often occur subsequently in self-compatible lineages (Charlesworth and Charlesworth 1979; Stone 2002; Takayama and Isogai 2005; Covey et al. 2010). Similarly in Brassicaceae, breakdown of SI appears to occur via loss of pistil-side factors, although the specific genetic mechanisms controlling SI are distinct between these two families (Kusaba et al. 2001; Sherman-Broyles et al. 2007; Tang et al. 2007; Shimizu et al. 2008). Several analyses in both these families have documented genome-wide effects of loss of SI (Hu et al. 2011; Slotte et al. 2013), however in most cases the associated transition to selfing has been recent. In contrast to these studies, all examined *Jaltomata* species have been found to be self-compatible (Mione 1992) (J. L. Kostyun and T. Mione, unpubl. data), indicating that the loss of self-incompatibility occurred early within *Jaltomata*, prior to the origin of the major subgroups of lineages within this clade (i.e. >3 MYA).

Consistent with an early loss of SI in this genus, by remapping RNA-seq reads (Wu et al. 2018) to the *J. sinuosa* genome we found that the heterozygosity level in all the 14 investigated *Jaltomata* species is comparable to that in the self-compatible species of *Solanum* but much reduced relative to that in the self-incompatible species of *Solanum* (Pease et al. 2016) (see Supplementary Information; Table S8). This relatively early loss of SI might also explain why we identified the significant contractions of gene families related to pollen-pistil interactions. A similar pattern was observed in *Caenorhabditis nigoni*, a self-fertile nematode species that diverged from its outcrossing sibling species *C. briggsae* approximately ∼3.5 MYA (Thomas et al. 2015). The *C. nigoni* genome revealed this self-fertile species has lost several hundred genes mediating protein-protein interactions, including the genes important for sperm-egg interactions (Yin et al. 2018).

Unlike the contraction of gene families potentially associated with relaxed selection, we also identified expansion of gene families that might be associated with the evolution of novel floral traits. In particular, we detected recent tandem duplications of one interesting candidate gene *SEUSS* (Figure 5). *SEUSS* is a transcription factor, which in *A. thaliana* interacts with *LEUNIG* within a transcriptional co-repressor complex to negatively regulate the expression of homeotic MADS-box gene *AGAMOUS* (Franks et al. 2002; Franks et al. 2006; Sridhar et al. 2006). Mutations in *SEUSS* cause ectopic and precocious expression of *AGAMOUS* mRNA, leading to partial homeotic transformation of floral organs in the outer two whorls (i.e. sepal and petal) (Franks et al. 2002). *LEUNIG* and *SEUSS* also have a more general role in lateral organ patterning and boundary formation in *Arabidopsis*, possibly through interactions with transcriptional factors in the *YABBY* and KNOX gene families (Stahle et al. 2009; Lee et al. 2014).

These functional roles are especially interesting with respect to the origin of floral diversity in *Jaltomata*, as the two derived corolla shapes (i.e. campanulate and tubular) are formed due to differential accelerated growth rates and variation in petal fusion (i.e. boundary formation) (Kostyun et al. 2017). Thus, the function of *SEUSS* is closely related to the mechanism generating the corolla shape diversity in *Jaltomata*. In general, duplication and divergence of floral identity genes appear to play an important role in the evolution of floral morphology in plants, such as the expansion of MIKC^c^-type MADS box genes (Hileman et al. 2006; Panchy et al. 2016). Although we do not yet have expression data for individual tissues (such as individual floral organs), we found that *SEUSS* copies were expressed only in the reproductive tissues of some lineages, and that species differ in the expression of these SEUSS copies. Interestingly, the SEUSS genes showed more limited expression in the orange-fruited-lineages (Table S15) with the derived floral traits (i.e. higher extent of corolla fusion), suggesting that down-regulation of some or all copies of this negative regulator of floral development might have contributed to the evolution of novel floral (corolla) morphs in this group. Future work with tissue-specific expression and function across different *Jaltomata* species with distinct corolla shapes will allow us to further assess whether *SEUSS* variants are strongly implicated in floral trait divergence across *Jaltomata*.

### B. Genes with patterns of adaptive molecular evolution

While many of the 58 genes that are inferred to be under positive selection specifically within the *Jaltomata* lineage (Table S16) are involved in general plant processes, some are from functional classes associated with the unique reproductive trait variation observed within this clade. For example, several candidates are associated with pollen-pistil interactions, including an S-ribonuclease binding protein that mediates the degradation of self-pollen (Sims and Ordanic 2001; Kao and Tsukamoto 2004). Novel changes in these proteins might have followed the early loss of self-incompatibility in this genus, perhaps as a result of reduced constraint on previous functions and the adoption of new roles under the altered reproductive environment of self-compatibility. Other potentially interesting candidates include an auxin response factor, a WRKY transcription factor, and a NAC transcription factor, all of which function in multiple plant developmental processes (Johnson and Lenhard 2011). Although the plant NAC gene family is quite large and its members play key roles in numerous developmental and stress response pathways, this putative candidate is especially interesting as NAM (a NAC protein) has been shown to function in organ boundary specification, including in flowers of closely related *Petunia* (Souer et al. 1996; Zhong et al. 2016), and development of the novel floral forms in *Jaltomata* appears to be specifically associated with altered floral organ boundaries (Kostyun et al. 2017).

### *The* Jaltomata *genome as a tool for future comparative analysis*

Overall, the generation of a high quality *Jaltomata* genome enabled us to evaluate genetic changes specific to this diverse lineage and across the Solanaceae more generally, as well as to clarify the historical evolutionary relationships among these important clades. In doing so, we inferred that *Jaltomata* has a complex history of divergence from its most closely related genera, which involves substantial introgression following initial lineage divergence. Based on quantification of recent TE activity, we infer that TE dynamics are likely an important contributor to genome content and genome size differences among groups in the Solanaceae. Using comparisons of gene family expansion and contraction, and more conventional analyses of adaptive protein evolution, we identified several genetic changes that might have either facilitated or accompanied the unique reproductive trait evolution observed in this clade.

In the future, the interesting evolutionary features and central phylogenetic placement of *Jaltomata* within the Solanaceae offers a unique opportunity to examine patterns of coding, structural, and transposable element evolution across this diverse and speciose plant family, as well as to further evaluate mechanisms that could contribute to the rapid reproductive diversification observed specifically within *Jaltomata*. These future comparative genomic analysis could reveal additional large and small scale genetic changes responsible for genomic differentiation and divergent form and function among a set of economically-important clades that are well-established models for developmental biology (*Petunia*), reproductive interactions (*Nicotiana*), secondary metabolite production (e.g., *Capsicum*), and ecophysiological responses (e.g., *Solanum*), as well as within this emerging model for analyzing the evolution of novel floral evolution.

## ACKNOWLEDGEMENTS

The authors thank David Haak for the flow cytometry estimate of *J. sinuosa* genome size, Shujun Ou for advice on transposable element analyses, and Matthew Hahn for advice on analyses and inference. This work was supported by the National Science Foundation [DEB 1135707 to L.C.M.].

## Supplementary Information

### Estimation of base-call error rate in the assembly

To obtain an upper bound on the error rate, we calculated the total base discrepancies between the primarily aligned individual Illumina reads and the assembly using Qualimap2 (Okonechnikov et al. 2016). To obtain a lower bound of error rate estimate, we mapped Illumina short reads back to the assembly by BWA v.0.7.12 (Li and Durbin 2010) and called variants with SAMtools v.0.1.19 (Li et al. 2009). Assembly error rates were estimated by dividing the number of detected variants (i.e. sites with a nucleotide consistently supported by Illumina reads but different from that in the assembled reference) and indels by the total length of covered genomic regions with mapping quality of more than 20 and mapped read depth ≥ 5.

### Estimating the heterozygosity level in *Jaltomata* species

To estimate the average heterozygosity in *J. sinuosa* and 13 other *Jaltomata* species that were previously investigated with transcriptome data (Wu et al. 2018), we mapped these RNA-seq reads from each of the 14 species to the *Jaltomata* genome assembly using STAR v2.5.2 (Dobin et al. 2013). SAM files generated were converted to sorted BAM files using SAMtools v. 0.1.19 (Li et al. 2009). SAMtools *mpileup* was then used to call alleles from the BAM files for all lineages, requiring non-reference allele calls to have Phred sore ≥ 30 and mapped read coverage ≥ 10. Heterozygosity was estimated by dividing the number of heterozygous sites by the total number of sites. Consistent with an early loss of SI in this genus, we found that the heterozygosity of all the 14 investigated *Jaltomata* species (Table S8) is comparable to that in the self-compatible species of *Solanum* but much reduced relative to that in the self-incompatible species of *Solanum* (Pease et al. 2016).

### Gene structure annotation

We used three different classes of evidence--RNA-seq data, protein homology, and *ab initio* gene prediction--in the MAKER2 pipeline v2.31.9 (Campbell et al. 2014) to annotate gene models. Assembled transcripts were generated from RNA-seq data sampled from multiple vegetative and reproductive tissues of the same accession of *J. sinuosa*, which are described in our previous phylogenomic study (Wu et al. 2018). Homologous protein sequences were downloaded from the SwissProt *Solanaceae* protein dataset (Magrane and Consortium 2011). For *ab initio* gene predictions, we used three programs SNAP (Korf 2004), GeneMark-ES (Lukashin and Borodovsky 1998) and AUGUSTUS v3.2.3 (Stanke et al. 2006). Each gene model was predicted using a two-pass (iterative) MAKER2 workflow. The initial SNAP HMMs were generated by CEGMA (Parra et al. 2007). After that, we ran the MAKER-P first round with the generated SNAP and GeneMark HMMs along with other evidence sets (i.e. transcriptome data and aligned *Solanaceae* protein set), while setting the parameters as “est2genome=1” and “protein2genome=1”. The gene predictors were then retrained one additional time, with the parameter setting as “est2genome=0” and “protein2genome=0”. The results from the MAKER-P *ab-initio* gene predictions were converted to the updated SNAP and AUGUSTUS HMMs, which were used for the second round of MAKER-P.

### Gene functional annotation

To assign gene functions, we first conducted sequence homology searches using BLASTP on the predicted protein sequences, against different protein datasets from SwissProt protein knowledgebase and its supplement TrEMBL (Magrane and Consortium 2011), Arabidopsis genome annotation TAIR10 (Lamesch et al. 2012), and tomato genome annotation ITAG3.2 (The Tomato Genome Consortium 2012). Putative protein domains on protein sequences were identified using InterProscan v5.25 (Jones et al. 2014). Finally, we used the pipeline AHRD (https://github.com/groupschoof/AHRD) to automatically select the most concise, informative and precise function annotation, with SwissProt, TAIR10, ITAG2.4, and TrEMBL being scored with different database weights (100, 50, 50, 10, respectively).

### Phylogenomic analyses

Four different approaches were used to perform phylogenetic reconstruction: 1) maximum-likelihood (ML) applied to concatenated alignments; 2) consensus of gene trees; 3) quartet-based gene tree reconciliation; and 4) Bayesian concordance of gene trees. The ML concatenation tree was inferred by using the GTRGAMMA model in RAxML v8.23 with 100 bootstraps (Stamatakis 2006). The consensus tree with internode certainty (IC) and tree certainty all (TCA) support scores were also generated using RAxML with the option for Majority Rule Extended (Salichos and Rokas 2013). The quartet-based estimation of the species tree was inferred by using the program ASTRAL v.4.10.9 with 100 bootstraps (Mirarab and Warnow 2015). Finally, the Bayesian primary concordance tree and associated concordance factors (CFs: IC and TCA) at each internode of the primary concordance tree was computed in the program BUCKy v1.4.4 (Larget et al. 2010). The input of a posterior distribution of gene trees was generated from an analysis with MrBayes v3.2 (Huelsenbeck and Ronquist 2001). We ran MrBayes for one million Markov chain Monte Carlo (MCMC) generations, and every 1000th tree was sampled. After discarding the first half of the 1000 resulting trees from MrBayes as burnin, BUCKy was performed for one million generations with the default prior probability (*α* = 1) (Larget et al. 2010). All inferred species trees were plotted using the R package “phytools” (Revell 2012).

### Phylogenetic analysis of genes in organelle genomes

We used the pipeline “Organelle_PBA” to reconstruct the chloroplast and mitochondrion genomes in *Jaltomata* using PacBio reads (Soorni et al. 2017). During the assembly, the chloroplast genome of *Solanum lycopersicum* (GI: 007898.3) and the mitochondria genome of *Nicotiana tabacum* (GI: 56806513) were used as the reference genome to filter PacBio sequencing data. Genes on the chloroplast and mitochondrion genomes were then annotated using DOGMA (Wyman et al. 2004) and Motify (Alverson et al. 2010), respectively. We downloaded the chloroplast and mitochondrial protein-coding gene sequences of *S. lycopersicum, C. annuum*, and *N. attenuata* from NCBI (Table S18). The orthologous genes were aligned using PRANK v.150803 (Löytynoja and Goldman 2005). The phylogenetic tree of chloroplast and mitochondrion were reconstructed with the concatenated sequences using maximum likelihood with the GTRGAMMA model in RAxML v8.23 (Stamatakis 2006).

### Differentiating introgression phylogeny from species tree

To differentiate which topology among *Jaltomata, Capsicum*, and *Solanum* most likely represented the initial pattern of lineage splitting (i.e. the ‘true’ species tree) versus subsequent introgression among species, we compared the relative divergence times (node depths) among species using gene trees that supported these two most conflicting bipartitions, with *Nicotiana attenuata* as the outgroup. In a rooted three-taxon phylogeny (i.e. “((P_1_, P_2_), P_3_), O” where P_1_, P_2_, and P_3_ are either *J. sinuosa, S. lycopersicum* or *C. annuum* and O is the outgroup *N. attenuata*), there are two divergence times: the earlier time (T_1_) when the first taxon P_3_ diverges from the remaining sister pair and the time T_2_ when the paired taxa P_1_ and P_2_ diverge (Fig. 3C, main text). We estimated T_1_ and T_2_ from biallelic informative sites, where allelic patterns can be represented as combinations of ancestral alleles (A) and derived alleles (B). T_2_ was calculated as: 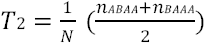 and T_1_ was calculated as 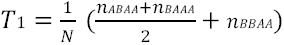, in which *N* is the number of sites (Fontaine et al. 2015). When P_3_ is the source of introgression, T_2_ is predicted to be lower which represents the time of introgression. When P_1_ or P_2_ is the source of introgression, both T_1_ and T_2_ will be lower. It is because that the true T_2_ will instead be the observed T_1_ and the time of ingression will be the observed T_2_.

### Examination of SEUSS gene copies within *Jaltomata*

Because our gene family analysis detected one important transcription factor locus *SEUSS* that showed rapid expansion specifically on the branch leading to *Jaltomata*, we further examined this apparent recent evolution of multiple copies of *SEUSS* genes in the *Jaltomata* genome, and their expression in different *Jaltomata* lineages. Using the tomato *SEUSS* gene in a homologous search, ten partial or complete copies of *SEUSS* (including seven copies annotated by the MAKER-P pipeline) were identified on two scaffolds (*scf29960* and *scf31961*) of *J. sinuosa* genome. Only a single copy was identified in each of the six other *Solanaceae* species. To confirm that the putative duplication events occurred after the split of Jaltomata from the other species/genera analyzed, we generated the gene tree of *SEUSS* among those *Solanaceae* species using maximum likelihood with the GTRGAMMA model in RAxML v8.23 (Stamatakis 2006), and showed that all inferred duplicate copies from Jaltomata were grouped within this tree (Fig. 5A, main text). We took several approaches to confirm the existence of multiple copies of *SEUSS* (i.e., to exclude assembly error). First, we searched for PacBio reads that spanned any two copies of identified *SEUSS* loci. Second, we examined the read depth of the Illumina short reads across the two relevant scaffolds to evaluate whether the read depth around each *SEUSS* copy was equal to or higher than read depth of the adjacent single-copy loci and genomic background. In cases where read depth is higher, this is likely due to an excess of multi-mapped reads at loci that have experienced very recent additional duplications that therefore cannot be distinguished during assembly. Third, using existing RNA-seq data from vegetative (seven tissues) or reproductive (four tissues) tissue pools among 13 different *Jaltomata* lineages (Wu et al. 2018), we examined whether each copy of the *SEUSS* locus was expressed or not in each tissue pool in each species. The program featureCounts (Liao et al. 2013) was used to assign either uniquely or multi-mapped reads to the genic regions of *SEUSS* from the generated input BAM files. Uniquely-mapped reads were used to distinguish the expression of specific *SEUSS* gene copies in each species, especially at structurally-intact *SEUSS* copies.

**Figure S1.**
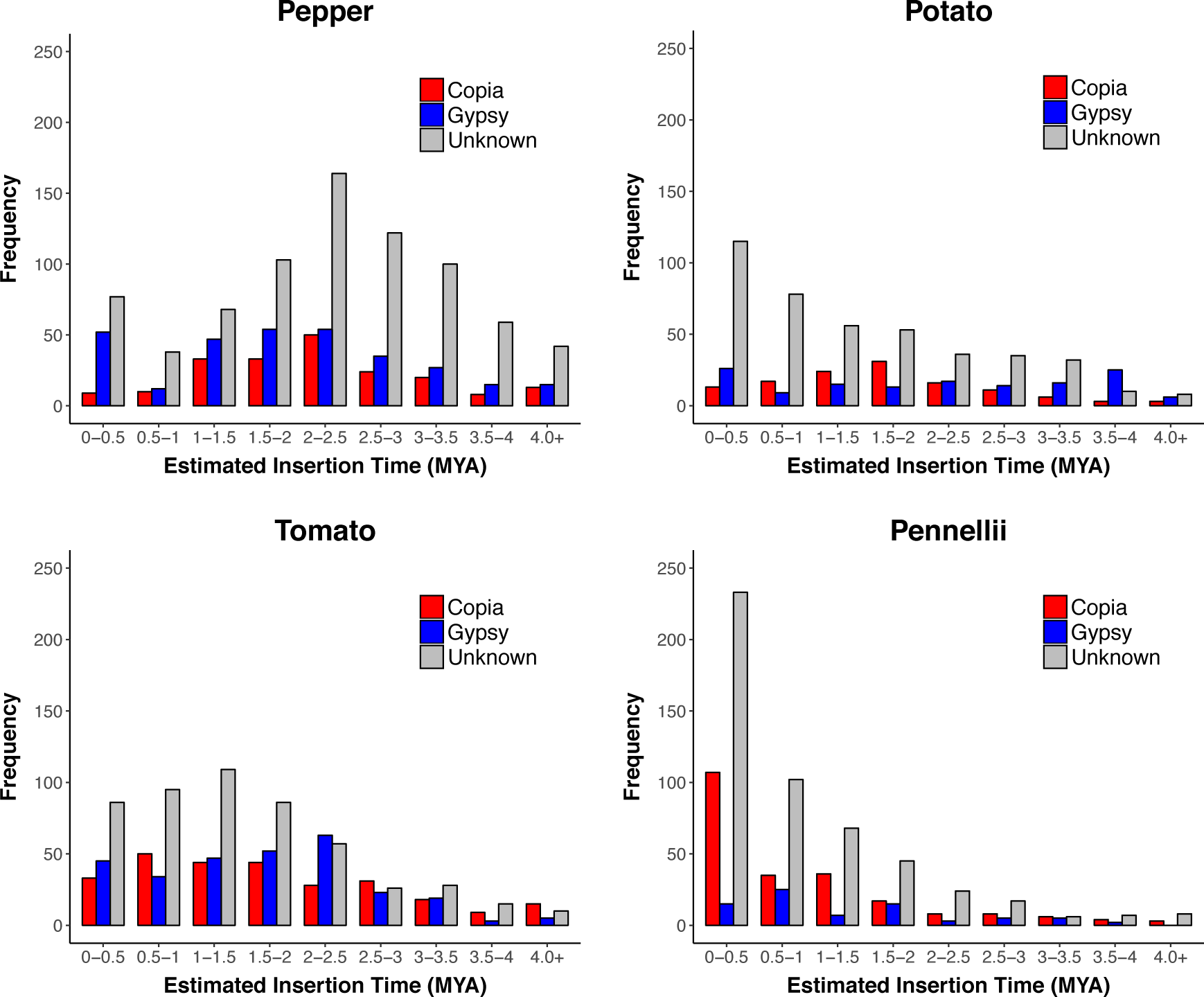
The estimated LTR-element insertion age distributions in **(A)** *C. annuum* (pepper), **(B)** *S. tuberosum* (potato), **(C)** *S. lycopersicum* (tomato), and **(D)** *S. pennellii*.

**Figure S2.**
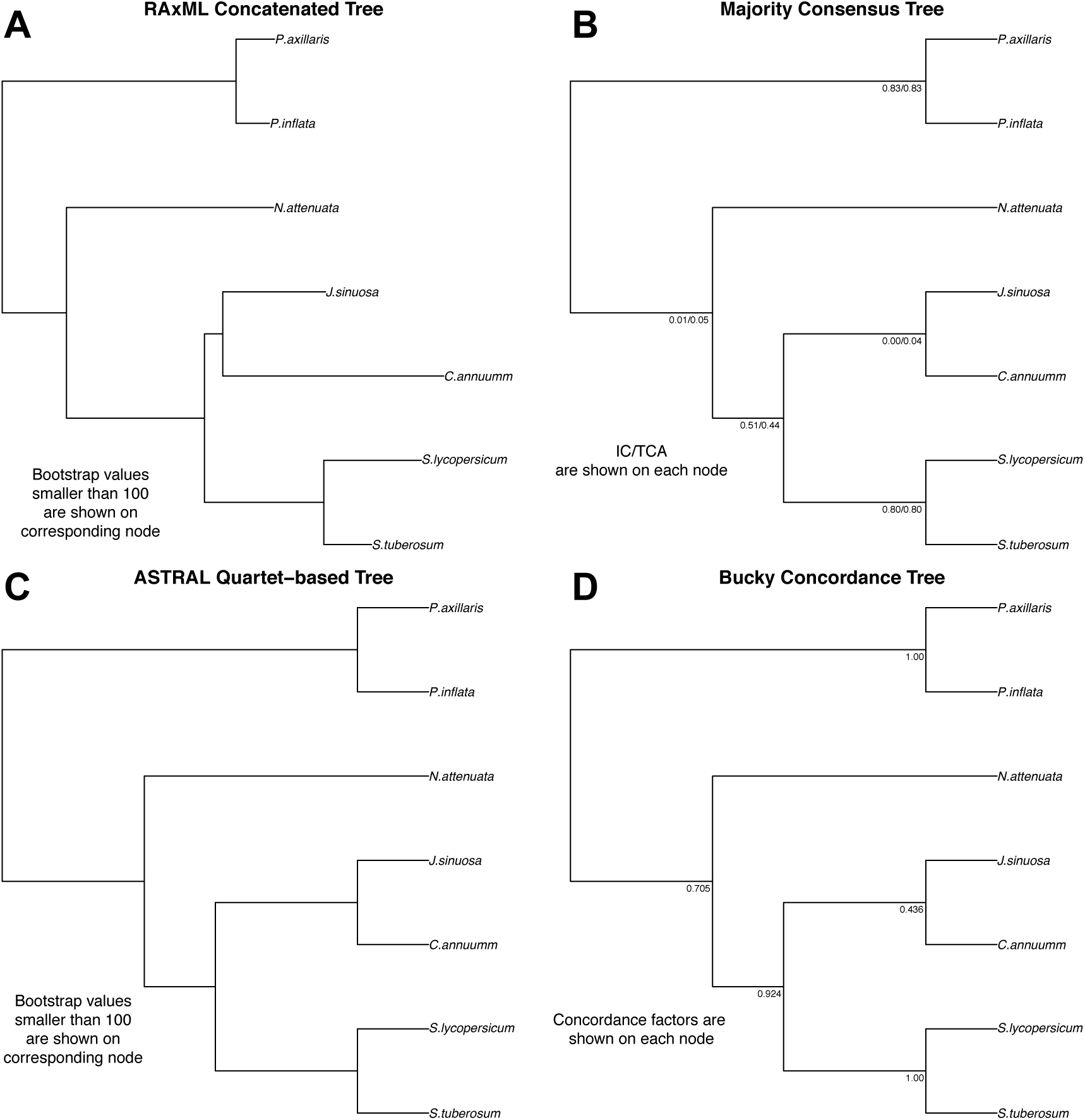
Whole-transcriptome phylogeny inferred by four different approaches on 3,103 single-copy 1-to-1 orthologs. **(A)** Whole-dataset concatenated phylogeny (RAxML). **(B)** Majority rule phylogeny (RAxML) with IC/TCA scores. **(C)** Best-likelihood quartet-based phylogeny (ASTRAL) inferred from 100 replicates. **(D)** Primary concordance phylogeny produced by BUCKy (α = 1) from gene trees inferred by MrBayes v3.2.1.

**Figure S3.**
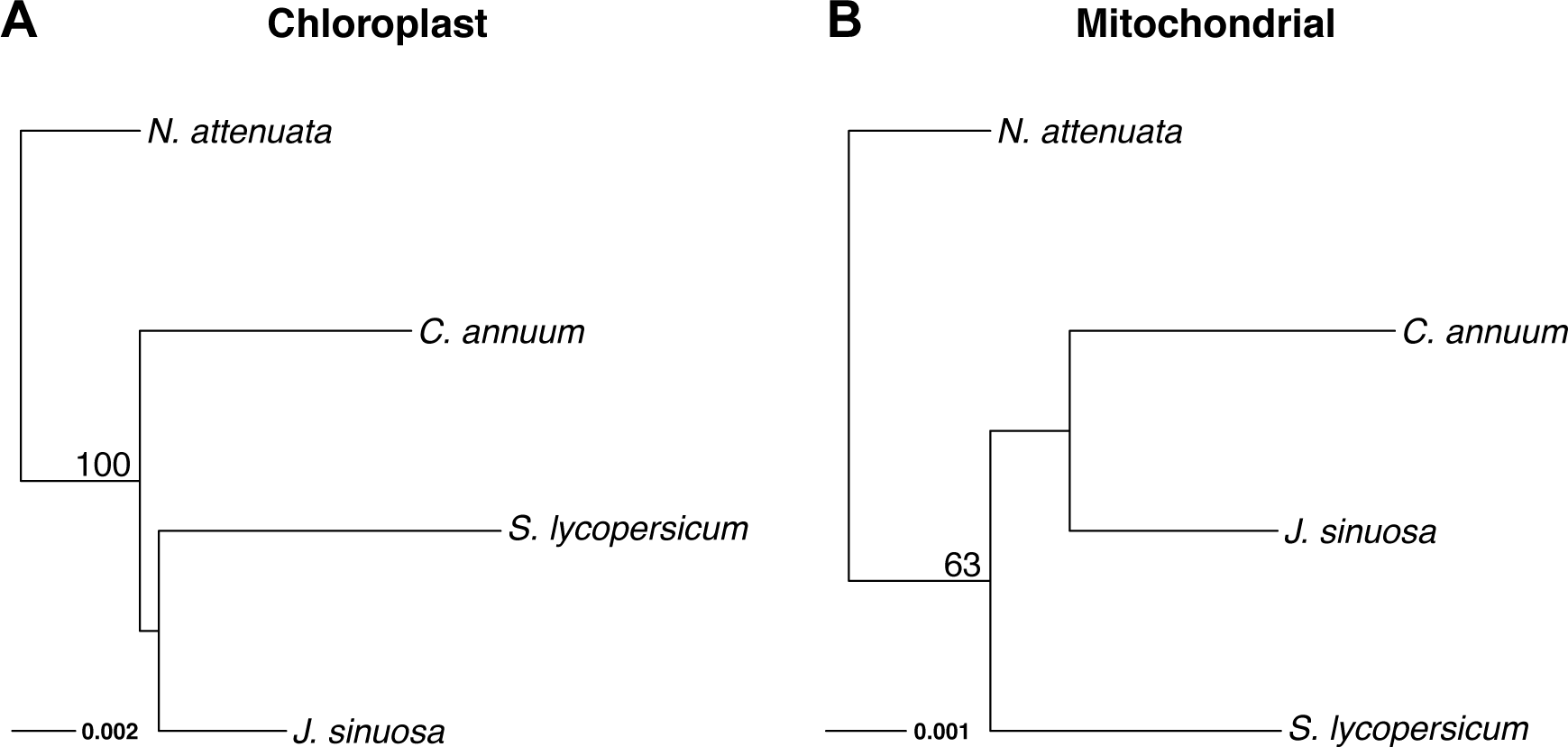
The concatenated phylogenies generated (in RAxML) using **(A)** 72 concatenated chloroplast genes and **(B)** ten concatenated mitochondrial genes. Bootstrap support is indicated above the focal branch.

